# Assessing Ebola Virus Circulation in the Tshuapa Province (Democratic Republic of the Congo): A One Health Investigation of Wildlife and Human Interactions

**DOI:** 10.1101/2024.09.17.613482

**Authors:** Rianne van Vredendaal, Léa Joffrin, Antea Paviotti, Claude Mande, Solange Milolo, Nicolas Laurent, Léa Fourchault, Douglas Adroabadrio, Pascal Baelo, Steve Ngoy, Papy Ansobi, Casimir Nebesse, Martine Peeters, Ahidjo Ayouba, Maeliss Champagne, Julie Bouillin, Jana Těšíková, Natalie Van Houtte, Sophie Gryseels, Maha Salloum, Freddy Bikioli, Séverine Thys, Jimmy Mpato, Ruben Ilonga, Henri Kimina, Ynke Larivière, Gwen Lemey, Pierre Van Damme, Jean-Pierre Van Geertruyden, Hypolite Muhindo-Mavoko, Patrick Mitashi, Herwig Leirs, Erik Verheyen, Guy-Crispin Gembu, Joachim Mariën

## Abstract

The wildlife reservoir and spillover mechanisms of the Ebola virus remain elusive despite extensive research efforts in endemic areas. This study employed a One Health approach to examine the virus’ circulation in wildlife and the associated human exposure risks in the Tshuapa province of the Democratic Republic of the Congo. In 2021, we collected 1015 samples from 888 animals, predominantly small mammals, and 380 inhabitants of Inkanamongo village, the site of an Ebola virus disease outbreak in 2014. These samples were screened for evidence of current (RNA) or past (IgG antibodies) Ebola virus infections. We also conducted interviews with 167 individuals in the surrounding districts to assess their interactions with wildlife. While no Ebola virus RNA was detected in the wildlife samples, anti-orthoebolavirus IgG antibodies were found in 13 bats and 38 rodents. Among the human participants, 120 individuals had IgG antibodies against at least one orthoebolavirus antigen, with 12 showing seropositivity for two antigens of the same orthoebolavirus, despite not having a prior Ebola disease diagnosis. Furthermore, over 50% of respondents reported frequent visits to the forest to hunt a variety of wild animals, particularly ungulates and rodents, which could account for occasional viral spillovers. The absence of active Ebola virus circulation in wildlife may reflect seasonal patterns in reservoir ecology, like those observed in bats. Similarly, seasonal human activities, like hunting and foraging, may result in periodic exposure risks. These findings highlight the importance of continuous, multidisciplinary surveillance to monitor changes in seasonal spillover risks.

## Introduction

Ebola disease is a severe viral illness characterised by haemorrhagic fever caused by viruses of the family *Filoviridae*, genus *Orthoebolavirus*. From 1976 to 2023, these viruses have led to 39 confirmed emergences across Africa [1]. The largest outbreak occurred between 2014 and 2016 in West Africa, where it caused over 28,000 reported cases and more than 11,000 deaths [2]. So far, six orthoebolaviruses have been identified: Ebola virus (EBOV, formerly known as Zaire ebolavirus), Sudan virus (SUDV), Bundibugyo virus (BDBV), Taï Forest virus (TAIFV), Reston virus (RESTV), and Bombali virus (BOMV). Among these, EBOV is the most lethal, with a fatality rate of up to 90% in humans in certain outbreaks [1]. It has also been responsible for most outbreaks, followed by SUDV and BDBV [3–7].

EBOV was first identified in 1976 in Yambuku, located in the Equateur province of the Democratic Republic of the Congo (DRC). Since then, EBOV has caused a total of 14 confirmed emergences in various regions of the DRC [1]. One occurred in 2014 in the Boende Health District (Tshuapa Province, western DRC), lasting from July 26 until October 7. This outbreak resulted in 69 reported cases (suspected, probable, and confirmed) and 49 deaths [8]. Most of these cases were reported in the Djera Sector, a forested area south of Boende. The outbreak’s index case was traced back to a pregnant woman from the village of Inkanamongo, who had butchered a monkey found dead by her husband [8].

Most outbreaks caused by EBOV, including the Inkanamongo outbreak, are believed to have a zoonotic origin, initiated by the virus spilling over from an animal to a human [8–10]. Nevertheless, despite extensive research efforts, the animal reservoir of the virus remains unidentified [10]. Over 12000 and 34000 wild animals have been screened respectively for EBOV RNA or antibodies between 1978 and 2023 to elucidate the reservoir and other animal hosts of EBOV (**Supplementary Table S1**) [5,10–16]. Gorillas, chimpanzees, and duikers have been suspected (and in the case of chimpanzees, confirmed) to have infected the index case during outbreaks in Gabon and the Republic of Congo (ROC) between 1996-2003 [17–20]. EBOV RNA has been detected in duikers, and both RNA and antibodies against EBOV have been found in great apes [19,21]. Although the presence of antibodies indicates previous exposure to the virus and detection of EBOV RNA proves infection at the time of sampling [17,22], their high disease-related mortality rate and limited distribution make these animals improbable reservoirs of EBOV [21,23].

Substantial evidence suggests that bats, or at least particular species, may serve as a reservoir for EBOV. Bats have been linked to the start of two different EBOV outbreaks in the DRC (in 2007, the index case had bought freshly killed fruit bats for consumption [24]) and in West Africa (the 2014-2016 outbreak likely originated from a spillover event involving a bat and a two-year-old boy [25]). Supporting this theory, EBOV antibodies were detected in at least eight frugivorous and two insectivorous Old World bat species [11,12,26–30]. Additionally, viral RNA has been found in three of these species (*Epomops franqueti*, *Hypsignathus monstrosus*, and *Myonycteris torquata*) [31]. Unlike apes or duikers, bats have not shown EBOV disease-related mortality, further indicating their potential role as a natural reservoir for the virus. Other filoviruses have also frequently been found in bats: *Rousettus aegyptiacus* is an important reservoir for the zoonotic Marburg virus, and Bombali, Lloviu and Měnglà viruses have so far only been detected in bats (respectively *Mops condylurus*, *Miniopterus schreibersii*, and *Rousettus sp*.) [32–35].

The notorious absence of detection of EBOV in wildlife or identification of the reservoir, despite extensive research effort, and in contrast to relatively frequent detections of other filoviruses in wild bats, could be due to several reasons. Possibly, EBOV only circulates in a limited number of species which have not been sampled sufficiently up to date. Considering the high mammalian diversity in Afrotropical forests, obtaining a sufficient sample size for each of the hundreds of species is unrealistic. Furthermore, EBOV infection dynamics could be highly seasonal, following the host reproductive cycles and life phases, and potential migrations [36–38], as has been observed for other mammalian-borne infections (e.g., paramyxoviruses, filoviruses, coronaviruses, arenaviruses, poxviruses, hantaviruses) [36,37,39–53]. The timing of many of the reservoir-searching studies (often conducted a few weeks after the start of an EBOV outbreak in humans) could have coincided with low incidence periods due to these seasonal changes in the natural epidemiology, making the detection probability too low.

Several studies have also investigated how socio-cultural factors and human behaviours contribute to the emergence of EVD outbreaks [54]. For instance, the social dimensions of EVD and viral haemorrhagic fevers were studied in Sierra Leone and Guinea [54,55]; bats and wild meat hunting and consumption and perceptions of disease risk were studied in Cameroon [56–58] and Ghana [59]. In the DRC, studies have linked a human EVD outbreak to exposure to fruit bats [24], highlighting that contact with bats, rodents, and eating non-human primate meat was associated with EBOV seropositivity, even in the absence of EVD diagnosis [60]. Additionally, multiple studies have investigated community beliefs about the origins of EVD [57,61,62].

Despite considerable efforts to identify the EBOV reservoir species and spillover pathways, extensive studies that simultaneously cover virological, ecological, sociological, and human health aspects are rare [63]. In line with the “One Health” concept, our study investigates EBOV circulation and health risks associated with wild meat hunting and consumption in a Congolese village (Inkanamongo, Boende Health District) where an Ebola outbreak occurred in 2014. Our study specifically aimed to (i) pinpoint the animal reservoir and ecological factors involved in EBOV maintenance and transmission, (ii) assess past human exposure to EBOV via a serological assay, and (iii) explore interactions between humans, wildlife, and the environment via sociological questionnaires. Although most of the confirmed Ebola disease emergences in the DRC have been caused by EBOV, we did not limit our study to EBOV but we also screened for other orthoebolaviruses. This study was conducted in the framework of the EBOVAC3 project, which ran a clinical trial (EBL2007) to assess the safety and immunogenicity of an Ebola vaccine in health care providers in Boende and aims to characterise outbreak preparedness through social science research.

## Methods

## 1. Animal reservoir study

### Trapping and sampling of wild mammals

Considering the assumed seasonality of EBOV outbreaks in the DRC, we conducted the ecological survey from May 13 to June 11, 2021, which coincides with the months preceding the 2014 outbreak (June-July). We trapped 633 small mammals (rodents, shrews, and bats) using a variety of traps within a 3 km radius of Inkanamongo (**Figure 1**). Additionally, we sampled 247 mammals provided by local inhabitants, including rodents captured in and around their homes and larger wildlife caught by hunters deeper in the forest. We refer to the **supplementary material** for more details on the sampling process.

**Figure 1:**
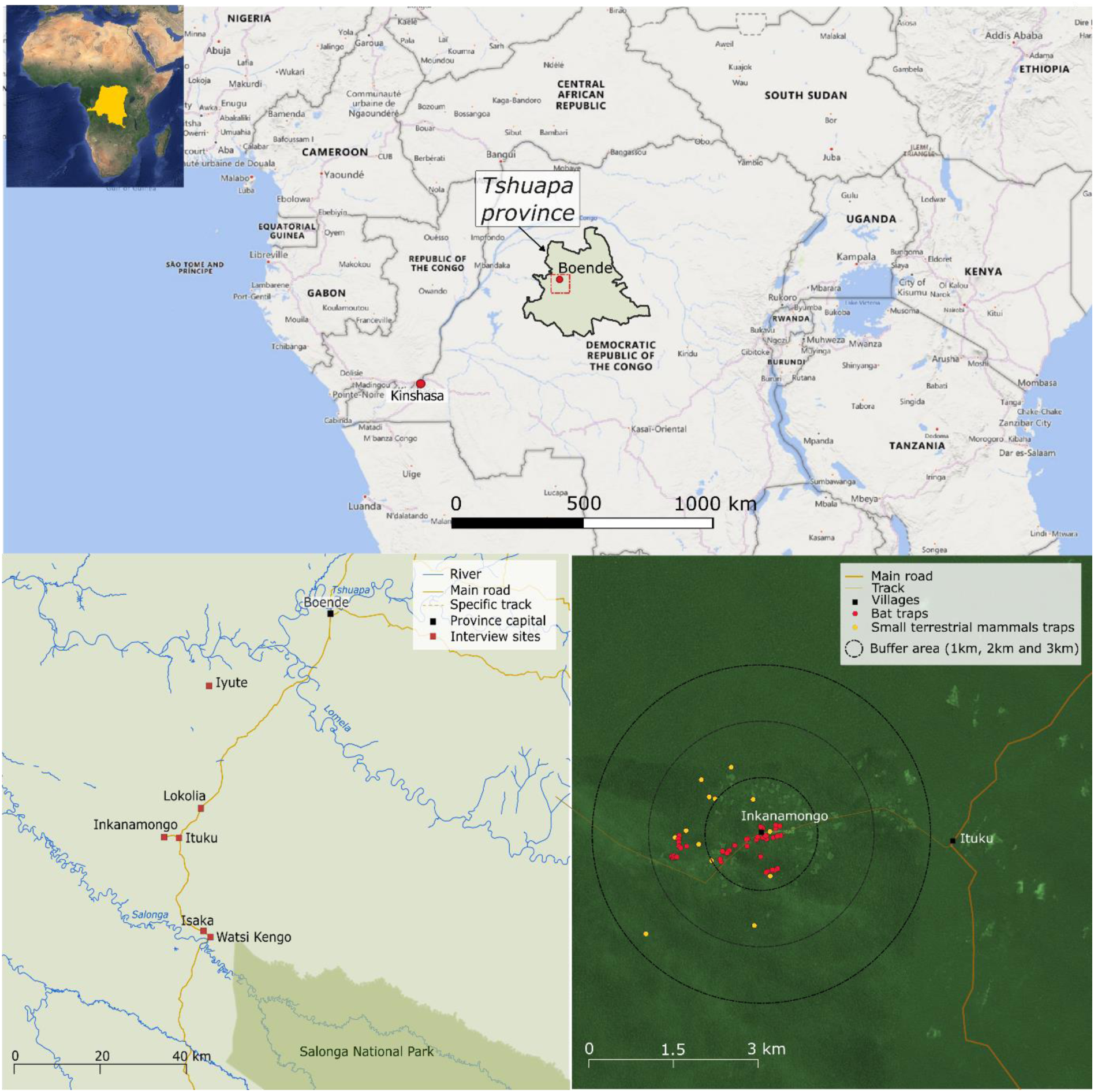
Sampling area, interview villages (Inkanamongo, Ituku, Lokolia, Iyute, Isaka, Watsi Kengo) in the Tshuapa province and trapping locations in and around Inkanamongo village.

We dissected all captured small terrestrial mammals, but bats were kept alive and only dissected in certain cases listed in the **supplementary material**. Live-captured terrestrial animals were euthanised by an overdose of isoflurane before the collection of blood on Whatman filter paper (hereafter called dried blood spots (DBS)) and tissue samples in DNA/RNA shield or 99% ethanol. For bats, we collected oral and rectal swabs, faeces, and urine (if available) stored in DNA/RNA Shield. We also performed a 3mm diameter wing punch biopsy for molecular identification and stored it in 99% ethanol. Whole blood was collected as DBS by puncturing the cephalic vein. All animals were preliminarily identified *in situ* to genus or species level using previously published identification keys [64–66] and pictures of each animal were taken in the field. Data regarding their sex, age, weight, and length were recorded.

### Molecular work

The inactivated samples in DNA/RNA shield were transported to the University of Antwerp and processed under a Biosafety cabinet class II for molecular species identification (via sequencing of the cytochrome b gene) and filovirus RNA detection. Based on prior studies indicating a higher likelihood of detecting filoviruses (Marburg virus and EBOV) in the liver and kidney [38,67–69], we conducted extractions from these specific organs. Filovirus RNA detection was carried out using two previously published PCR assays with degenerate primers targeting a fragment of the filovirus polymerase (L) gene [70,71]. Full details of the procedures are provided in the **supplementary material** and in **Table S2**.

### Animal Serology

We used a multiplex serological assay (Luminex-based technology) to test DBS from bats and rodents for anti-orthoebolavirus IgG antibodies, as previously described [11,14,15]. In brief, the assay incorporated recombinant orthoebolavirus proteins, including glycoprotein (GP), nucleoprotein (NP), and viral protein 40 (VP40) from five different orthoebolaviruses (EBOV, SUDV, BDBV, RESTV, and BOMV). To identify a sample as positive, we used various combinations of antigens for which the median fluorescence intensity (MFI) exceeded the estimated cutoffs. Combinations with more than one antigen of the same orthoebolavirus above the cutoff values suggest that a sample is more likely to be truly positive. All details are provided in the **supplementary material**.

In the absence of established wildlife control samples, we determined cut-off values using several methods tailored to the available data. For bats, we calculated the mean MFI cut-off values based on four different approaches. This included the mean MFI plus 4 times the standard deviation from 145 negative samples obtained from bats housed in European zoos [11], as well as the 97.5th percentile cut-off derived from three statistical analyses (change point, exponential, and binomial) using a larger dataset of wild bat samples (totalling 8,741 individuals from Guinea, DRC, and Cameroon) [14].

For the rodent samples, due to the lack of validated negative control samples and a comprehensive panel of previously tested wildlife specimens, we calculated cut-offs directly from our study samples. Specifically, the MFI cut-off was set at the 97.5th percentile, recognising that this method inherently produces at least a 2.5% false positive rate. To mitigate this, we only classified a sample as positive if both the nucleocapsid and glycoprotein of the same orthoebolavirus exceeded their respective cut-offs. Because both antigens are not correlated in false-positive samples, this criterion significantly reduces the chances of false-positive results. We performed the analyses with R version 4.3.0 software [72].

## 2. Human serology study

A cross-sectional study was conducted in Inkanamongo in the Boende Health District and the Lokolia Health Area where there was an Ebola virus disease (EVD) outbreak in 2014. The door-to-door survey was conducted from December 2 to 5, 2023, to estimate seroprevalence against orthoebolaviruses in the human population with a sample size of 380 adults who had never received a vaccine against Ebola virus. Blood samples were collected on Whatman 903 filter paper from each participant’s fingertips. Each participant provided information on socio-demographic characteristics, knowledge of and information channels about EVD, whether they had experienced symptoms of the disease, and whether they or a close relative had contracted EVD in the past. Samples were tested for anti-orthoebolavirus antibodies using the same Luminex assay as previously described for wild animals, but with positivity-cutoffs calculated on human positive and negative control samples as explained in detail in [73]. To identify participant characteristics associated with past EBOV infection, we developed different (Generalized) Linear Models for each EBOV antigen with IgG titres (expressed as the logarithm of MFI values) or antibody presence (higher or lower than cutoff) as response variable and age, sex, and previous EVD (no, yes, or indirect contact only via family member) as explanatory variables [74]. These models were developed for each EBOV antigen. We conducted our analyses using R version 4.2.2 software [72]. The significance of the explanatory variables was evaluated using χ2 tests (p-values) or by assessing changes in the Akaike Information Criterion (ΔAIC), which involved comparing the full model against models with individual variables removed.

## 3. Social science study

We conducted two types of social science studies: interview sessions in Inkanamongo (referred to as Study A) and a more structured study conducted in villages of the entire Djera Sector (referred to as Study B). **Study A** corresponds to observations and interview sessions conducted in Inkanamongo between May 17 and May 27, 2021, simultaneously with the animal samplings. This study consisted of two sub-surveys. During the first one, observation notes were taken about the inhabitants’ daily activities. Following a convenience sampling method, 40 inhabitants were interviewed with the help of a structured questionnaire to assess (i) their daily activities and (ii) their behaviours regarding animal capture, specifically assessing local hunting behaviours and preferences. The second sub-survey focused specifically on residents’ (i) awareness, perceptions, and knowledge of bats, (ii) hunting habits regarding bats and (iii) beliefs or practices associated with bats. Ten persons were interviewed with the help of a structured questionnaire. Additional information on study A is provided in the **supplementary material**.

**Study B** refers to a dedicated survey mission conducted in six villages of the Djera Sector most affected by Ebola in 2014 (Inkanamongo, Lokolia, Ituku, Watsi Kengo, Isaka, Iyute; **Figure 1**). Here, we interviewed 117 inhabitants of these villages to understand the potential of animal-to-human and human-to-human transmission of EBOV by assessing (i) the people’s interactions with the forest by investigating their use and knowledge of the environment (agriculture, fishing, and hunting); and (ii) their interactions between and within human communities. Data was collected door-to-door between July 16 and July 23, 2022. All details are provided in the **supplementary material** and **Table S3**.

## Results

## 1. Animal reservoir study

We collected samples from 888 animals (**Table 1**), of which 18% of bats (70 out of 382) and 35% of rodents (107 out of 306) were opportunistically collected from hunters. Overall, we did not detect any active EBOV infection in 1015 screened samples (430 pooled non-invasive samples and 585 pooled kidney and liver). A substantial proportion (23%) of female bats were either late pregnant or lactating, specifically 39.1% of *E. franqueti* (54 out of 138), 39.3% of *M. torquata* (11 out of 28), and 40% of *H. monstrosus* (4 out of 10).

**Table 1.**
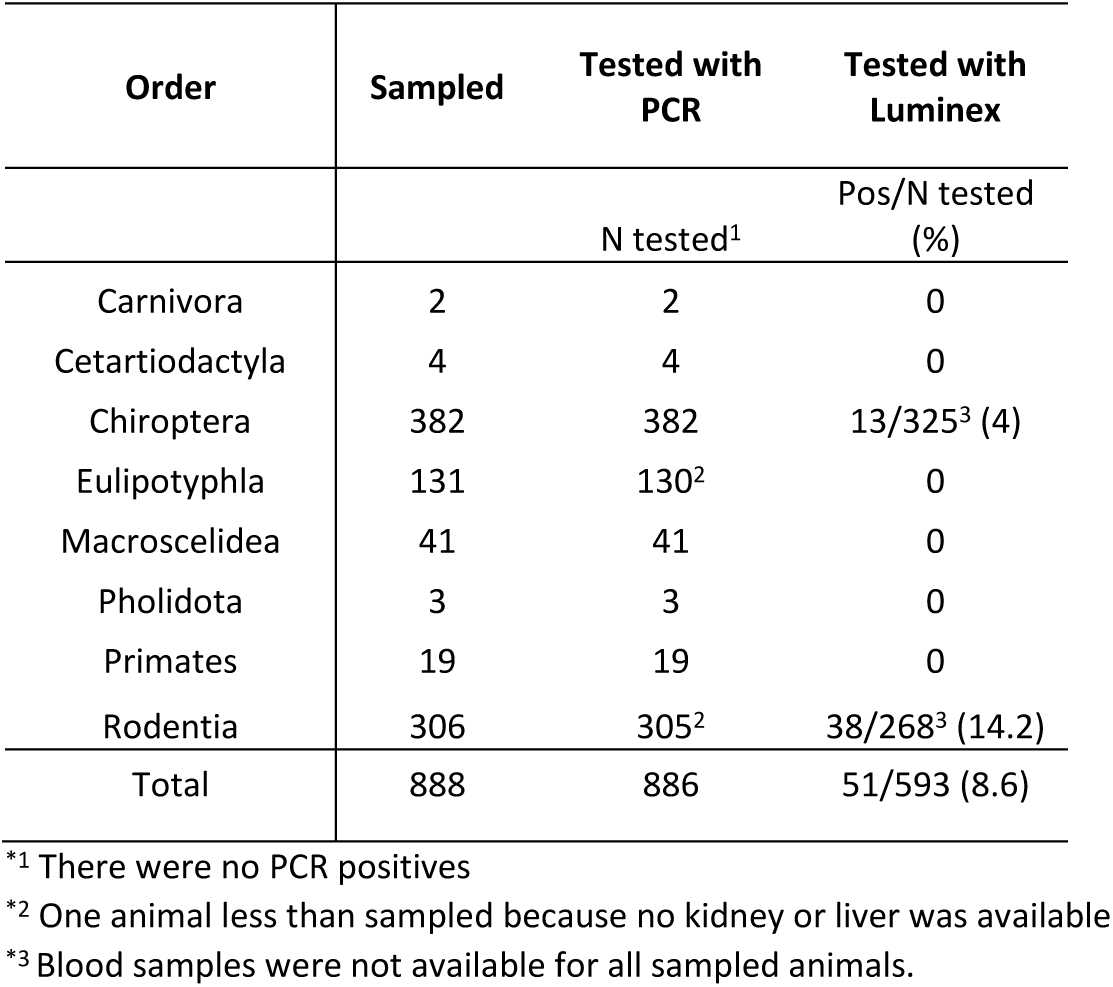
Overview of the number of animals sampled, tested with PCR, and tested with Luminex per taxonomic order. The number of animals reactive with at least one orthoebolavirus antigen and the percentage are presented.

Among the 325 bats screened, 13 were reactive with at least one orthoebolavirus antigen (**Table 2**). All bats that showed reactivity belonged to the Pteropodidae family, encompassing four distinct species: *H. monstrosus* (2/13), *E. franqueti* (3/13), *Eidolon helvum* (2 /13), and *M. torquata* (6/13). Cross-reactivity among the same antigens of different orthoebolaviruses was observed, with two bats testing positive for the GP antigens of EBOV, SUDV, and BDBV, and one bat testing positive for the VP40 antigens of EBOV, SUDV, and BDBV. However, no bats tested positive for combinations of different antigens (e.g., NP+GP or GP+VP40) from the same orthoebolavirus.

**Table 2.** Number and percentage of animals and humans that tested antibody positive against different orthoebolavirus antigens using a Luminex assay, based on two different cut-off values (for animals). The assay used recombinant proteins of Nucleoprotein (NP), Glycoprotein (GP), or Viral Protein-40 (VP40) for different orthoebolaviruses: Ebola virus (EBOV), Sudan virus (SUDV), Bundibugyo virus (BDBV), Reston virus (RESTV), and Bombali virus (BOMV). GP proteins from the Mayinga (GP-M) and the Kissidougou (GP-K) strain were used for EBOV.

| Taxon | Antigen | EBOV |  |  |  | SUDV |  |  | BDBV |  | RESTV | BOMV |
| --- | --- | --- | --- | --- | --- | --- | --- | --- | --- | --- | --- | --- |
|  |  | NP | GP-K | GP-M | VP40 | NP | GP | VP40 | GP | VP40 | GP | GP |
| Chiroptera<br>N=325 | Mean+4SD<sup>1</sup> | 0 (0) | 2 (0.5) | 1 (0.2) | 1 (0.2) | 2 (0.5) | 7 (1.4) | 3 (0.6) | 3 (0.6) | 1 (0.2) | 1 (0.2) | - |
|  | Mean 3 cutoffs<sup>2</sup> | 0 (0) | 1 (0.2) | 1 (0.2) | 1 (0.2) | 0 (0) | 1 (0.2) | 2 (0.5) | 0 (0) | 1 (0.2) | 1 (0.2) | - |
| Rodentia<br>N=268 | Mean+4SD<sup>3</sup> | 3 (1.1) | 5 (1.9) | 5 (1.9) | 1 (0.3) | 1 (0.3) | 5 (1.9) | 1 (0.3) | 5 (1.9) | 2 (0.7) | 3 (1.1) | - |
|  | Mean 3 cutoffs<sup>3</sup> | 7 (2.7) | 16 (6.2) | 19 (7.3) | 2 (0.7) | 4 (1.5) | 18 (6.9) | 1 (0.3) | 5 (1.9) | 1 (0.3) | 3 (1.1) | - |
| Humans<br>N=380 | Mean 3 cutoffs<sup>4</sup> | 8 (2.1) | 23 (6.1) | 42 (11.3) | 19 (5.0) | 14 (3.7) | 84 (22.1) | 2 (0.5) | 10 (2.6) | 14 (3.7) | 2 (0.5) | 0 (0.0) |

<sup>\*1</sup> calculated on a panel of 145 negative samples (bats) [11]
<sup>\*2</sup> calculated on a panel of 8741 wildlife samples (bats) [14]
<sup>\*3</sup> calculated on 268 samples from this study (rodents)
<sup>\*4</sup> calculated on a panel of 202 human samples [73]

From the 268 rodents tested, 38 rodents from nine different genera were reactive with at least one orthoebolavirus antigen (**Table 2**): *Funisciurus* (2/38), *Grammomys* (2/38), *Hylomyscus* (1/38), *Lophuromys* (15/38), *Mus* (3/38), *Oenomys* (1/38), *Praomys* (4/38), *Rattus* (9/38), and *Thamnomys* (1/38). However, given that the changepoint analyses suggested a considerably higher number of positives than the mean+4SD method (37 vs. 17), we assume that cutoffs might be too low and many samples could be false positives. Nevertheless, three samples are more likely to be true positives given that two different antigens from the same orthoebolavirus scored positive: one *Lophuromys* sp. tested positive for a combination of VP40+GP EBOV antigens, one *Lophuromys* sp. for a combination of the NP and GP EBOV antigens, and one *Lophuromys* sp. showed reactivity against the NP and GP antigens of SUDV. Most of the rodents that showed reactivity against at least one orthoebolavirus antigen were captured in or around the village (24/38); one of these was the *Lophuromys* sp. showing simultaneous reactivity for NP and GP EBOV antigens.

## 2. Human serology study

We screened DBS from 380 participants between 18 and 76 years old and found IgG antibodies against at least one orthoebolavirus antigen in 120 participants (31.6%, 95% CI: 26.9-36.5%. **Figure 2**, **Table 2**). Reactivity to at least two antigens of the same orthoebolavirus (much stronger indication for true positive samples) was observed for 12 of 380 individuals (3.2%, 95% CI: 1.6-5.5%), of which seven reacted to EBOV antigens, three to SUDV, one to BDBV, and one to both EBOV and BDBV (**Table 3; Figure S1**). Of these 12 seropositive individuals, ten were male and two were female, and their ages ranged from 18 to 52 years. Interestingly, none of these 12 individuals were previously diagnosed with EVD themselves, although two of them reported they had someone close to them who suffered from EVD (**Table S4**). In contrast, out of eight individuals who had contracted EVD in the past, only two were found to be reactive with one antigen (GP EBOV and GP SUDV), and none showed reactivity against two antigens. The (generalised) linear models did not suggest any significant effects of the age, sex or previous EVD status on the presence or absence of antibodies or their titres against EBOV (**Table 4**).

**Figure 2.**
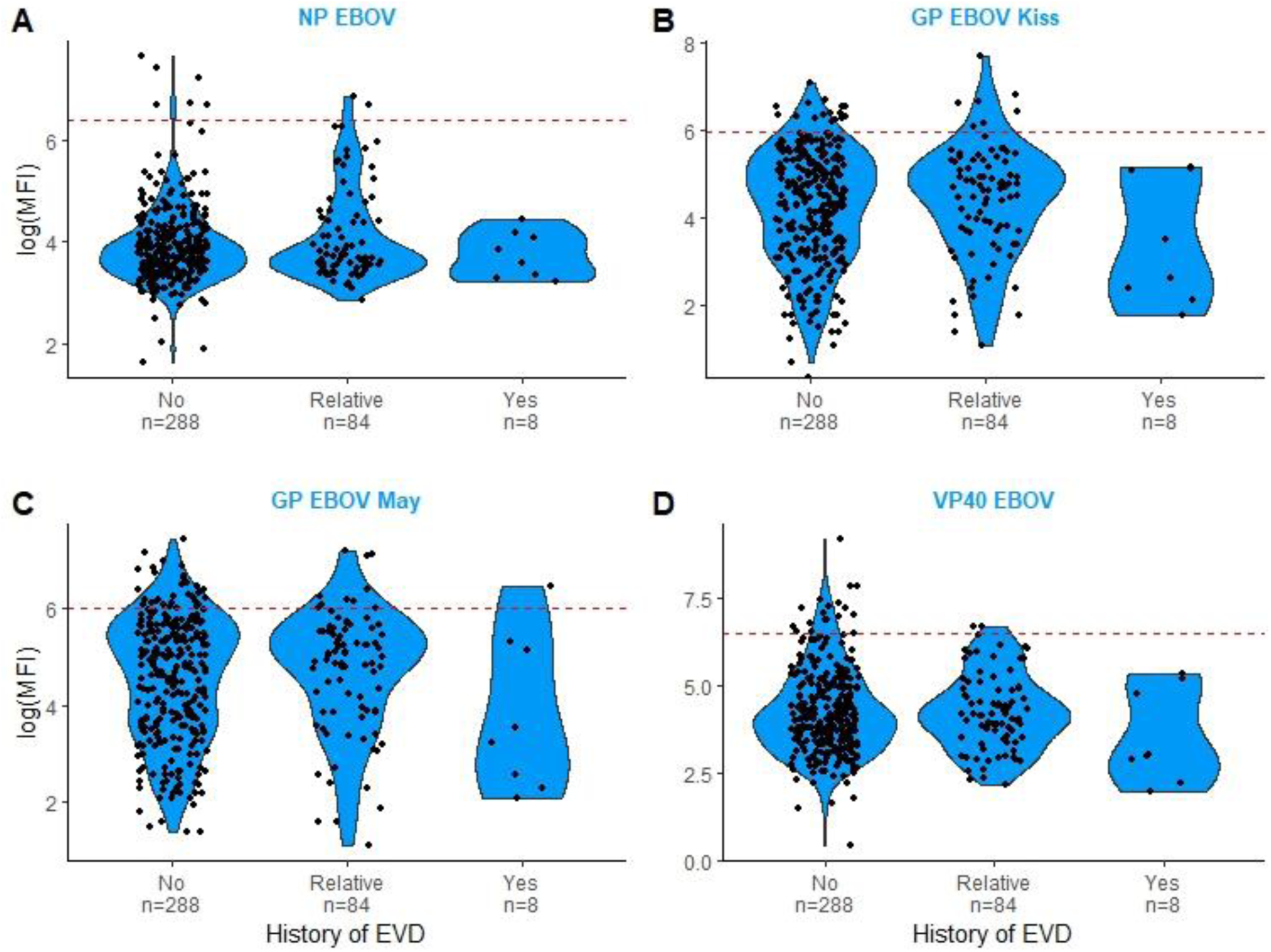
IgG antibody titres against Ebola virus from inhabitants in Inkanamongo (DR Congo), expressed as Median Fluorescent Intensity values (MFI) on the Luminex for four different antigens: nucleocapsid protein (NP), the glycoprotein Kissidougou (GP-Kiss), the glycoprotein Mayaro (GP-Kiss) and the viral protein40 (VP40). *Violin plots express the log(MFI) values for three different categories of participants according to previous Ebola viral disease (EVD) exposure: no infection, relative became infected, and the participant became infected. The red dotted line indicates the cutoff of the antigen on the Luminex. Positivity cutoffs calculated on a panel of 202 human positive and negative control samples as explained in detail in* [73].

**Table 3.** Number and percentage of samples reactive with at least two antigens of the same orthoebolavirus out of 380 total samples from humans.

| Virus | EBOV |  |  |  | SUDV |  | BDBV |
| --- | --- | --- | --- | --- | --- | --- | --- |
| Antigens | NP + GP-K + GP-M + VP40 | NP + GP-K + GP-M | NP + GP-M | GP-K + GP-M + VP40 | NP + GP | GP + VP40 | GP + VP40 |
| n (%) | 1 (0.3) | 3 (0.8) | 3 (0.8) | 1 (0.3) | 1 (0.3) | 2 (0.5) | 2 (0.5) |
| n (%) per orthoebolavirus | 8 (2.1) |  |  |  | 3 (0.8) |  | 2 (0.5) |

**Table 4.** Overview of the different models used to test the effect of EVD, age, and sex on MFI and seropositivity in humans for each antigen.

| Response | Antigen | NP |  | GP-Kiss |  | GP-May |  | VP40 |  |
| --- | --- | --- | --- | --- | --- | --- | --- | --- | --- |
|  | Test stat | Pval | ΔAIC | Pval | ΔAIC | Pval | ΔAIC | Pval | ΔAIC |
| MFI (LM) | EVD | 0.11 | 0.03 | 0.10 | 0.10 | 0.12 | -0.21 | 0.27 | -0.92 |
|  | Age | 0.12 | -0.23 | 0.05 | 1.35 | 0.05 | 1.29 | 0.35 | -1.12 |
|  | Sex | 0.45 | -1.28 | 0.94 | -1.98 | 0.84 | -1.95 | 0.98 | -1.99 |
| Seropositivity (GLM) | EVD | 0.54 | -2.73 | 0.33 | -2.18 | 0.92 | -3.89 | 0.21 | -1.21 |
|  | Age | 0.42 | -1.61 | 0.12 | 0.07 | 0.22 | -0.67 | 0.15 | -0.46 |
|  | Sex | 0.6 | 1.48 | 0.81 | -1.95 | 0.42 | 1.38 | 0.99 | -1.99 |

## 3. Social science study

In the first sub-survey of study A, 40 people were interviewed explicitly about their livelihoods in the rural village of Inkanamongo. All interviewees belong to the Mongo ethnic group. Of the 40 interviewees, 77.5% (n=31) reported having two to three occupations. Thirteen respondents identified hunting as their main activity, one focused on fishing, 20 on farming, five were teachers, and one was a clerk. Overall, 32 people reported engaging in hunting, 28 in fishing, 25 in farming, five as teachers, one as a clerk, and one identified as a notable person. All participants stated that wild meat was primarily for local consumption, while 17 indicated that they also exported it to Boende. When asked about changes in the forest over recent years, 65% of respondents (n=26) reported that wild animals had become rare, 25% (n=10) observed a decline in soil quality and crop production, and 32% (n=13) mentioned a decrease in unspecified natural resources. One respondent did not report any particular change. Regarding the perceived causes of these changes, 87.5% of individuals (n=35) attributed them to overexploitation of the forest, while five pointed to a shift away from ancestral customs and practices that previously helped regulate resources. Additionally, seven linked the changes to overpopulation in the village, five believed it was God’s will, three attributed them to climate change, and one did not provide any specific cause.

Specific interviews about bats were conducted with ten adults (comprising seven men and three women) who represented six distinct Mongo ethnic clans: Ekolie, Djwangenda, Djiyoko, Nongo, Djeefo, and Djosona. Identifications of bats varied among respondents: two linked them to birds, five to dogs, and one to rabbits, with one describing them as “birds with dog teeth”. Regardless of ethnic clan, only men reported being involved in trapping bats. All the participants (n=10) could identify fruit bats as “Lolema” and insectivorous bats as “Ifuki.” Eight participants knew that the diet of bats consists of fruits or insects. Although only one respondent could indicate specific roosting sites for fruit bats (about 20 km from Inkanamongo on the banks of the Salonga River), all knew that bats often roost under house roofs (n=8) or trees (n=6). Interestingly, one person reported using the presence of bats as an indicator of wild fruit ripening, which he uses to plan the timing of fruit harvests. Spiritual beliefs related to bats were particularly reported about the Ekolie clan, where three people mentioned that vocalisations signal misfortune. For this reason, Ekolie only engages in hunting and selling bats, not in consuming them. Health risks associated with bats were largely disbelieved (n=4) or uncertain (n=5), except for one respondent who mentioned Ebola transmission via contact.

In Study B, we surveyed 117 individuals across six villages, enabling us to generalise the findings for the entire district. Wild meat is consumed by almost all respondents (n=109, 7 non-responses) and is typically (but not exclusively) prepared by women. Nearly all participants (n=113, with four non-response) reported engaging in agriculture (n=106), 57% engaged in fishing (n=65), and 50% in hunting (n=56). Although hunting was the primary occupation for only five respondents, many engaged in it as a secondary (n=30) or tertiary activity (n=21). Hunting typically occurred on ancestral lands, suggesting limited cooperation between villages in this activity. The frequency of forest visits varied, with nearly half of the respondents (n=54) entering daily and one-third (n=37) entering two to three times per week. Sixty-eight respondents noted that the frequency of forest visits increased during certain months, particularly in July, which marks the start of the “caterpillar season” and the availability of ripened fruits in the forest. This period also coincides with a reported increase in animal sightings, likely due to the greater abundance of food.

When combining data from the two surveys (A and B), most respondents (n=129) reported encountering a variety of animals in the forest (in total from 13 taxonomic orders), with the Cetartiodactyla (antelopes, sitatunga, bushpig) and Rodentia (porcupines, squirrels, rats) representing the orders most frequently reported as observed (respectively 34.5% and 29.8%) and reported as hunted (respectively 46.5% and 26.5%) (**Figure 3**). Only nine out of 13 orders observed in the forest were reported as hunted by the respondents. Molecular identification of wild meat samples collected in Inkanamongo confirmed the presence of most of the animals mentioned by hunters in the sociological study. Overall, we identified at least seven orders, 25 genera, and 27 species of animals hunted around Inkanamongo (**Table S5**). Interestingly, the Chiroptera order (bats) was not reported as hunted by any respondent. Nevertheless, during the animal reservoir study, we observed hunters returning to the village with dead bats. Similarly, we observed people in the village with dead elephant shrews (Macroscelidea order), but none of the respondents reported having observed or hunted them. Last, gorillas, chimpanzees, and lions were reported as observed by respondents, although they do not occur in the area. We provide in the **supplementary material** (**Table S6**) a comprehensive list of animals reported as present in the study area, as well as those for which a translation of the local names used during the interviews was available in the literature.

**Figure 3.**
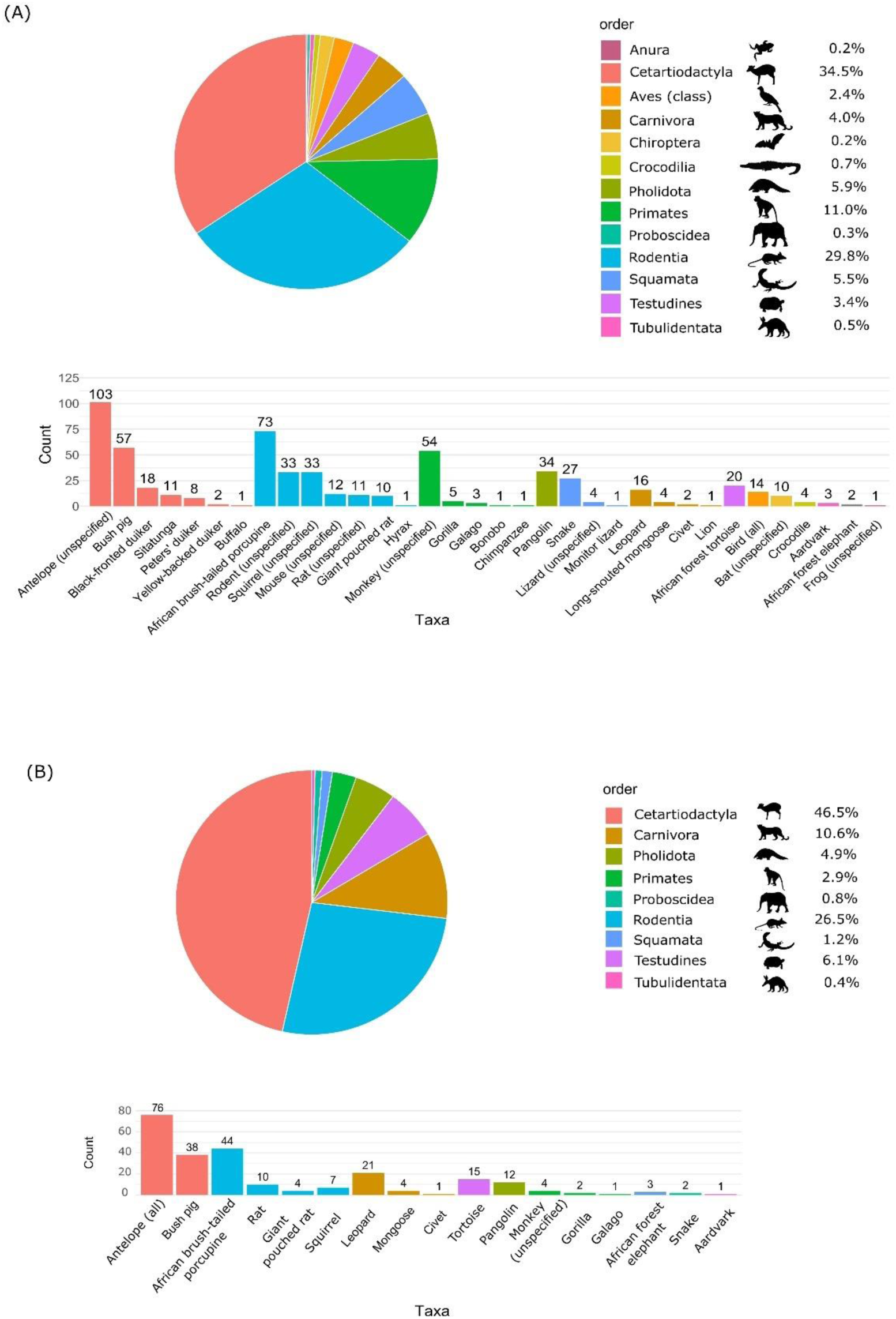
Mentions of different vertebrate taxa reported as seen in the forest (A; N=580 occurrences, all villages including Inkanamongo) and mentioned as hunted (B; N=245 occurrences, all villages including Inkanamongo). *Thirty mentions could not be assigned to any taxa or order as no translation could be found. We provide in the **supplementary material** a table with the translation of the different taxa when available (**Table S6**)*.

## Discussion

The detection of EBOV in wildlife remains challenging, as highlighted by the limited number of studies and the few species in which the virus has been detected [17,22,31]. Notably, all studies conducted in the DRC, both during and after EBOV outbreak periods, including ours, have been unsuccessful in detecting EBOV RNA in wildlife.

Although we did not detect any active EBOV circulation in wildlife around Inkanamongo, we identified antibodies against at least one EBOV antigen in some animals, indicating past exposure to the virus or to related orthoebolaviruses. Interestingly, the seroprevalence in our bat samples was lower compared to other studies conducted in the DRC and other endemic regions [12,14,15]. For instance, Lacroix et al. (2021) reported EBOV antibodies in 1.2% to 10.8% of bats in the DRC, depending on the positivity criteria [14]. Using the same methodology and acceptance criteria as Lacroix et al., we found antibodies in only 1.2% to 4.0% of bats. Similar to our findings, Lacroix et al.’s samples generally did not show simultaneous reactivity to different EBOV antigens, except for one *Epomophorus* bat from Nord Kivu province. Another serological survey conducted in 2023 across Western and Central Africa using the same methodology found even higher seroprevalence estimates in bats, ranging from 6.1% to 18.9% [15]. We assume that the observed differences in seroprevalence may be attributed to varying ecological factors, such as seasonality, inducing changes in reproductive status and migration patterns.

As nearly a quarter of the female bats in our study were either late pregnant or lactating, the reproductive status might have influenced our overall seroprevalence estimates. Afrotropical bats are reported in the literature to experience one or two reproductive seasons per year depending on the species, often synchronised with the rainy seasons to ensure ample food resources for their young [75,76]. Pregnant and lactating bats often experience altered immunological responses due to increased nutritional stress, changes in roosting behaviour, hormonal shifts, and varying energy allocation [77–81]. While studies on related viruses have found higher seroprevalence in pregnant and lactating bats due to the additional metabolic demands of these reproductive stages [79,80,82], previous research on EBOV infection in gestating or lactating bats has yielded mixed results. For example, gestating *Hypsignathus monstrosus* and *Lissonycteris angolensis* showed lower seroprevalence for GP-SUDV, while higher seroprevalence was observed in *Eidolon helvum* and *Micropteropus pusillus,* and no significant difference in seroprevalence was found in *Rousettus aegyptiacus* [15,83]. Although caution is warranted when interpreting these findings due to limitations of data aggregating from different sites, it seems that a seasonally fluctuating reproductive status could affect seroprevalence estimates in bat populations [15,83].

The lower seroprevalence estimates in the bat population around Inkanamongo may also be explained by the periodic migration of certain bat species during the sampling period [10,24]. Specifically, we observed the arrival of seasonally migrating *Eidolon helvum* bats from May to June. While no *E. helvum* were captured during the first two weeks of sampling in May 2021, we began capturing them in week three and observed hunters bringing them to the village. An interviewee also noted that the arrival of fruit bats signals the ripening of certain fruits, confirming the seasonal fluctuation of fruit bats around the village. While seasonal migration of bats offers increased hunting opportunities for the inhabitants, it may also result in certain times of the year with elevated risks for zoonotic disease transmission. Notably, many of the bat species in which we found EBOV antibodies – including *E. helvum*, *H. monstrosus*, *E. franqueti*, and *M. torquata* - were observed being brought to the village by hunters. This aligns with previous findings where EBOV RNA was detected in three of these species, suggesting their potential role as EBOV reservoirs in wildlife [31].

The interpretation of our serological findings in rodents requires caution due to the absence of validated negative control samples, which can influence the determination of a biased cut-off value. Nonetheless, three rodents tested positive for two antigens of the same orthoebolavirus, meeting a more stringent serological criterion [11,14]. Interestingly, all were *Lophuromys*, of which one was captured in the village. Additionally, traces of EBOV RNA were previously detected in three rodent taxa (*Mus setulosus* and two unidentified *Praomys species*) and one shrew (*Sylvisorex ollula*), although antigen detection and attempts to isolate the virus failed [84]. Furthermore, several studies have shown that filovirus-like elements are integrated into the genomes of certain rodents, shrews, and bats, indicating a long-term history of mammal-filovirus association [85,86]. Modelling has also suggested bats, rodents, and shrews as primary candidates for EBOV reservoirs [87,88]. However, as conclusive empirical evidence is still lacking, increased surveillance, accounting for potential seasonal variations, is needed to elucidate the role of rodents and shrews in EBOV circulation.

Another objective of our study was to assess human exposure to EBOV in Inkanamongo. While almost one-third of the inhabitants (n=120) had IgG antibodies against at least one orthoebolavirus antigen, we identified only 12 seropositive participants who were positive for at least two antigens of the same orthoebolavirus. Surprisingly, none of these 12 seropositive participants had a history of EVD. This finding is consistent with several other studies in which they detected EBOV antibodies in individuals with previously undetected EBOV infections and no EVD-related symptoms [5,55,74,89]. Although some of our participants reported experiencing EVD-like symptoms such as fever and vomiting in the past months, these symptoms could also be attributed to other diseases, and we cannot exclude the possibility of false-positive results. Nevertheless, our results indicate that the inhabitants of Inkanamongo are occasionally exposed to orthoebolaviruses in their surroundings and/or that EBOV may present as a minimally symptomatic or asymptomatic infection. Additionally, we observed that none of the previously EVD-infected survivors in the villages had antibodies against at least two EBOV antigens. This indicates that humoral immunity may decline more rapidly than anticipated, a phenomenon also observed in survivors of the 2013-2016 EBOV outbreak in Sierra Leone [90] and Guinea [91].

Finally, we investigated the interactions between humans and wildlife in the villages. Our findings indicate that over 50% of the humans in the area frequently visit the forest to hunt various wild animals, particularly ungulates and rodents, which poses a considerable risk of exposure to zoonotic diseases. However, we noticed inconsistencies between the respondents’ reported behaviour and our field observations concerning human-wildlife interactions. We observed inhabitants with dead elephant shrews although they did not report hunting them, they may have misclassified them as rodents. Additionally, although no respondents mentioned hunting bats in the region, we documented instances where some people engaged in selling, hunting, or consuming bats in Inkanamongo. Also, specific interviews on bats revealed that ethnic-based perceptions and tradition are important factors to consider when investigating interactions with bats. Such sociocultural aspects have been partially investigated in the context of EVD emergence [92]. However, epidemiologic links between wildlife consumption or various aspects of practices and infections remain poorly understood. The discrepancy between interviews and field observations may indicate differences in interests directly related to wildlife hunting (e.g., livelihoods or trade) or awareness of health risks, which may lead to reluctance among individuals to admit hunting and consuming bats, particularly following the Ebola outbreak in 2014. Disentangling the health risk factors and socioeconomic dynamics of disease emergence remains a complex issue and requires further interdisciplinary investigation in the future [93].

Furthermore, during our questionnaire administrations, significant uncertainties emerged about the origin of the 2014 Ebola outbreak in Inkanamongo. While Maganga et al. (2014) reported that the outbreak started on the 26^th^ of July 2014 in Inkanamongo, the head nurse at Lokolia Health Centre (a pivotal figure in the local health system) stated that the index case originated in Tomoge, a village approximately 12 kilometres from Lokolia. He mentioned that the index in this village (the wife of a teacher) discovered a dead antelope (*Tragelaphus eurycerus* or bongo) in the forest and consumed it with her family. This index case became ill on July 15^th^, nine days preceding the documented start of the outbreak by Maganga et al. (2014) Interestingly, the individual considered to be the official index case from Inkanamongo was the 28^th^ person to be recorded with Ebola-like symptoms, according to the Lokolia Health Centre records. According to the head nurse, this person fell ill after visiting an infected woman in Inkanamongo, who had contracted it from another village. This alternative version of the outbreak is coherent with the testimony of the husband of the presumed index case that we interviewed in Inkanamongo. This questioning of the outbreak narrative, years after the initial investigation into the outbreak’s origin, highlights the challenges of epidemiological traceability. Accurate identification of the index is crucial for eco-epidemiological studies and subsequent field missions aimed at monitoring host animals and zoonotic diseases. Misidentifying the index case may lead to incorrect geographical areas and potentially to the wrong suspected reservoir animals of the outbreak.

### Conclusion

Despite not detecting active EBOV infection in our samples, our serological results indicate that inhabitants and wildlife in this region are occasionally exposed to EBOV or other orthoebolaviruses. These findings align with our sociological study, which revealed that most people frequently enter the forest to hunt various wildlife that are suggested to be potential EBOV reservoirs, including rodents, bats, antelopes, and primates. Unfortunately, the broad range of wildlife hunted by respondents did not allow us to pinpoint the most likely EBOV reservoir. We hypothesise that the lack of active EBOV infection during our study may be due to seasonal patterns in the migration or reproductive cycles of the presumed reservoir host. Additionally, we noticed that changes in human activities, such as increased hunting and foraging during periods of food abundance (e.g., caterpillars, fruits, animals), could periodically elevate the risk of wildlife disease spillover. Therefore, we recommend year-round surveillance to monitor seasonal variations in the presence of EBOV or other zoonotic diseases in endemic villages.

## Acknowledgments

We wish to thank Loic Adrien Mbong Osseke for his contribution to the molecular identification of potential reservoir species.

## Author Statement

The following contributions are listed in alphabetical order: <u>Conceived the study</u>: Guy-Crispin Gembu, Herwig Leirs, Joachim Mariën, Patrick Mitashi, Hypolite Muhindo-Mavoko, Antea Paviotti, Martine Peeters, Séverine Thys, Pierre Van Damme, Jean-Pierre Van Geertruyden, Erik Verheyen. <u>Data collection (animal reservoir data)</u>: Douglas Adroabadrio, Papy Ansobi, Pascal Baelo, Guy-Crispin Gembu, Léa Joffrin, Herwig Leirs, Nicolas Laurent, Claude Mande, Joachim Mariën, Casimir Nebesse, Steve Ngoy, Erik Verheyen, Rianne van Vredendaal. <u>Data collection</u> <u>(sociological data)</u>: Papy Ansobi, Freddy Bikioli, Ruben Ilonga, Henri Kimina, Claude Mande, Hypolite Muhindo-Mavoko, Jimmy Mpato, Patrick Mitashi, Antea Paviotti, Maha Salloum, Séverine Thys. <u>Provided the human serological data</u>: Papy Ansobi, Solange Milolo, Hypolite Muhindo-Mavoko, Patrick Mitashi. <u>Performed the laboratory work (animal reservoir data)</u>: Ahidjo Ayouba, Pascal Baelo, Julie Bouillin, Maeliss Champagne, Léa Fourchault, Sophie Gryseels, Léa Joffrin, Nicolas Laurent, Claude Mande, Jana Těšíková, Natalie Van Houtte, Rianne van Vredendaal. <u>Performed the laboratory work</u> <u>(human serological data)</u>: Julie Bouillin, Maeliss Champagne, Solange Milolo. <u>Performed the data</u> <u>analysis</u>: Léa Joffrin, Joachim Mariën, Antea Paviotti, Rianne van Vredendaal. <u>Supervision and</u> <u>coordination work (animal reservoir study)</u>: Guy-Crispin Gembu, Sophie Gryseels, Herwig Leirs, Joachim Mariën, Martine Peeters, Erik Verheyen<u>. Supervision and coordination work (sociological</u> <u>study)</u>: Ynke Larivière, Gwen Lemey, Antea Paviotti, Séverine Thys, Pierre Van Damme, Jean-Pierre Van Geertruyden. <u>Supervision and coordination work (human serology study)</u>: Ahidjo Ayouba, Patrick Mitashi, Hypolite Muhindo-Mavoko. <u>Wrote the paper</u>: Léa Joffrin, Joachim Mariën, Antea Paviotti, Rianne van Vredendaal. All authors read and approved the final manuscript.

## Availability of data and materials

The datasets used and/or analysed during the current study are available from the corresponding author upon reasonable request.

## Ethical clearance and permits

Ethical clearance for the sociological studies was approved by the Public Health Ethics Committee of the University of Kinshasa and provided by DRC’s National Committee for Health Ethics (N°368/CNES/BN/PMMF/2022, approved on 5/07/2022). Ethical clearance for the animal reservoir study was provided by the Ethical Committee for Animal Testing of the University of Antwerp (N°2020-22, approved on 03/04/2021). The Nagoya permit for the export of samples from the animal reservoir study was issued by the Ministry of Environment and Sustainable Development, General Secretariat, in Kinshasa (N°009/ANCCB-RDC/SG-EDD/BTB/09/2020 [Ministère de l’Environnement et Développement Durable, Secrétariat Général à l’Environnement et Développement Durable – in French]). The CITES permits for the animal reservoir study were provided by CITES/DRC Management Authority in Kinshasa (N°CDFF1918R, CDFF1919R, CDFF1920R, CDFF1921R [Organe de gestion CITES/RDC – in French]) and the FPS Health, Food Chain Safety and Environment, DG Environment, Multilateral Affairs and Strategic Matters, CITES Management Body, Brussels, Belgium (N°2022/BE00308/PI [FOD Volksgezondheid, Veiligheid Voedselketen en Leefmilieu, DG Leefmilieu, Dienst Multilaterale en Strategische Zaken, Beheersorgaan CITES - in Dutch]). All “movement of personal” (called “ordre de Mission”) were signed and approved by the local authorities at each site. Material transfer agreements (MTA) were issued by the University of Kisangani, Biodiversity Surveillance Center (CSB [Centre de Surveillance de la Biodiversité] – in French). Live captured animals were euthanised (isoflurane) following the 2013 AVMA Guidelines for the Euthanasia of Animals and Sikes and Gannon 2007 (J Mammal. 88:809–23).

