## Supplementary material for "Assessing Ebola Virus Circulation in the Tshuapa Province (Democratic Republic of the Congo): A One Health Investigation of Wildlife and Human Interactions"

### 1. Context

A substantial number of animals have been captured, sampled, and screened between 1978 and 2023 in the DRC to detect the EBOV animal reservoir (**Table S1**) [1–8]. Interestingly, the studies conducted a few weeks after the start of an EBOV outbreak in humans did not seem to have higher detection rates.

Biological evidence supports this stance: EBOV antibodies were detected in at least eight frugivorous and two insectivorous Old World bat species [3,4,9–14], and viral RNA in three of these species (*Epomops franqueti*, *Hypsignathus monstrosus*, and *Myonycteris torquata*) [15]. Additionally, certain bat species (*Mops condylurus, Chaerephon pumilus, and Epomophorus wahlbergi*) are, to our knowledge, the only mammals that can survive inoculation of EBOV and replicate the virus without showing signs of the disease [16]. Faecal shedding of the virus has been detected in *E. wahlbergi*, indicating a possible transmission route [16]. Experimental inoculation with EBOV has also been performed on *Rousettus aegyptiacus*, in which individuals showed antibody development but no or infrequent viral detection and no evidence of viral shedding [17,18]. Additionally, *R. aegyptiacus* is the established natural reservoir of a closely related filovirus, Marburg virus (MARV) [19], and *M. condylurus* bats are the suspected reservoir of BOMV [20–23]. This indicates that bats play an important role in the transmission ecology of (at least certain) filoviruses.

Also, outbreaks of MARV in humans have been linked to these risk seasons [24]. Detection rates of active MARV infection are high when targeting its reservoir *R. aegyptiacus*, especially during the birthing seasons. Considering the close relatedness between MARV and EBOV, a similar circulation pattern in bat populations could be expected for EBOV.

### 2. Material and methods

The study was conducted in Inkanamongo, a village in the Boende Health District with approximately 750 inhabitants. The village is surrounded by forest and only accessible via narrow forest paths. Crops like cassava and rice are cultivated on several crop fields located in the forest, both close to the village and deeper in the forest. Livestock, like pigs, sheep, and goats, as well as chickens, are free-roaming in the village. Most houses are made of wood or bricks, and the roof of straw. Inkanamongo's economy primarily relies on selling locally brewed alcohol and wild meat.

**Sampling details- Animal reservoir study**

Small terrestrial mammals were captured using a combination of Sherman live, snap, and pitfall traps to maximise the diversity. We placed 20 pitfall traps along a transect at 5-meter intervals, alternating with Sherman or snap traps positioned 5 meters away from each side of the transect. This combination of 60 traps on three transects was laid out in 9 different sites around the village. The sites were in primary forest, secondary forest, and old fallow land. Traps were left in the field for 7-10 days in each site, resulting in 4680 trap nights. A total of 40 Sherman live traps and snap traps were also placed in and around the houses in the villages from May 13 to June 9. Eighty-six squirrel traps were deployed in trees in the secondary forest for 27 nights. Traps were checked every morning and rebaited with a mixture of palm nut pulp, nuts, and fish, depending on availability. Bats were captured with mist nets and harp traps placed in flight corridors, next to fruit trees, and above streams in the primary and secondary forests and in young fallow land around the village. Between three to six mist nets and, at most, two harp traps were placed for 21 nights. Bats were rehydrated when removed from traps, after process, and before release; water and sugar were used for fruit bats, only water for insectivorous ones.

Bats were dissected (using the same methods as for rodents and shrews) only if they were found dead, unintentionally died during the collection or sampling process, were potentially from a rare or new species and needed to be preserved as voucher specimens, or if insufficient blood could be collected without endangering their lives (particularly in the case of small-sized individuals). Opportunistic samples were taken from bats that local hunters unsolicited provided. Additionally, we took tongue samples in 99% ethanol and rectal, oral, nasal, and urogenital swabs in DNA/RNA Shield for 56 carcasses of large mammals (pangolins, monkeys, etc.).

**Viral screening**

We extracted RNA from pooled kidney and liver samples of 584 individuals with the Macherey-Nagel Nucleospin RNA kit. Approximately 20 mg of tissue was used per extraction. RNA extractions of the pooled swab samples (combined with urine and faeces if available) of 370 bats were carried out using the Qiagen QIAamp Viral RNA mini kit. Swabs, faecal, and urine samples were briefly vortexed, centrifuged (1500 x g for 15min), and pooled per individual to extract faecal RNA using the Qiagen QIAamp viral RNA kit (Qiagen, Valencia, CA, USA) following the manufacturer's recommendations. RNA extraction of swabs taken from carcasses of 53 individuals hunted for wild meat was also done using the Qiagen QIAamp Viral RNA mini kit using 70µL of each of the four different swabs per individual (rectal, oral, nasal, and urogenital). Extraction was performed according to the manufacturer's instructions, but double elution using 2 x 40µL RNase-free $H_2O$. Reverse transcription was performed on 8 µL of RNA extract using Maxima Reverse Transcriptase (Thermo Scientific). Synthesised cDNA was screened for filoviruses using two PCR assays with degenerate primers targeting a fragment of the filovirus polymerase (L) gene (**Table S2**).

Amplification conditions for the first system (using GoTaq from Promega) included 94 °C for 2 min, followed by 40 cycles of denaturation (94 °C, 1 min), annealing (52 °C, 1 min), and elongation (72 °C, 1 min), and a final extension step of 72 °C for 5 min. The program for the first and second rounds of the nested PCR system (also using GoTaq from Promega) was denaturing at 94 °C for 2 min, followed by 35 and 30 cycles (respectively) of 94 °C denaturing for 30 s, 57 °C and 54 °C (respectively) annealing for 30 s, 72 °C extending for 40 s with final extension 72 °C for 5 min. The amplification products were analysed by electrophoresis in 1.5% agarose gels stained with 0.5 µg/mL GelRed Nucleic Acid Stain in TBE (Tris/Borate/EDTA) buffer. Visualisation occurred under UV light.

**Molecular species identification**

For samples extracted with the Qiagen Qi

Molecular species identification was done using the RNA extracts of samples extracted with either the Qiagen QIAamp kit or the Macherey-Nagel kit as described above. DNA was extracted from the samples specified above using the Qiagen QIAamp 96 DNA QIAcube HT kit according to manufacturer's instructions, apart from performing sample disruption with Zymo bashing beads for 15 s on max speed on a Bertin Minilys. Next, cytochrome b PCR was performed using primers L14723 (5'-ACCAATGACATGAAAAATCATCGTT-3') and either H15915 (5'-TCTCCATTTCTGGTTTACAAGAC-3') or H15149 (5'-GCCCCTCAGAATGATATTTGTCCTCA-3'). Amplification conditions (using GoTaq from Promega or Platinum Taq II from ThermoFisher Scientific) included 94 °C for 5 min, followed by 40 cycles of denaturation (94 °C, 30 s), annealing (52 °C, 30 s) and elongation (72 °C, 90 s), and a final extension step of 72 °C for 10 min. Positive PCR amplicons were either purified using the ExoSAP-IT protocol, following the manufacturer's instructions and sequenced by Macrogen Europe (Netherlands) or were purified and Sanger sequenced at Neuromics Support Facility (Antwerp, Belgium). All mitochondrial raw sequences were trimmed and assembled using Geneious Prime (Biomatters Ltd., Auckland, New Zealand). The consensus sequences were aligned with ClustalW, MAFFT or MUSCLE (as implemented in Geneious Prime) to control for gaps, translate the sequences into amino acids, map the primer pairs, and trim the consensus sequences.

**Animal Serology**

The assay incorporated recombinant orthoebolavirus proteins, including glycoprotein (GP), nucleoprotein (NP) and viral protein 40 (VP40) from 5 different orthoebolaviruses: EBOV (NP, amino acids [aa] 488 to 739, variant Mayinga 1976; VP40, aa 31 to 326, variant Kissidougou-Makona 2014; GP-K, aa 1 to 650, variant Kissidougou-Makona 2014; GP-M, aa 1 to 650, variant Mayinga 1976); SUDV (NP, aa 361-738, variant Gulu; GP, aa 1 to 637, variant Uganda 2000; and VP40, aa 31 to 326, variant Gulu); BDBV (GP, aa 1 to 501, variant Uganda 2007; VP40, aa 31 to 326, variant Uganda 2007); RESTV (GP, aa 1 to 650) and BOMV (GP, aa 1 to 672). Following sample dilution (corresponding to a final plasma dilution of 1/2000), we incubated 100 µL with 50 µL of magnetic beads coated with recombinant protein (2 µg protein/1.25 x $10^6$ beads) and washed hereafter. For bats, we added 0.1 µg/mL of goat anti-bat biotin-labeled IgG (Euromedex, Souffelweyersheim, France) to each well and incubated for 30 min at 400 rpm at room temperature. For rodents, we added biotin anti-mouse IgG (4µg/mL)208 (Sigma-Aldrich B7022; Merck Life Science; Hoeilaart, Belgium) to each sample. Shrews and elephant shrews were not analysed due to the unavailability of suitable secondary antibodies. After a new washing step, we added 50 µL of 4 µg/mL streptavidin-R-phycoerythrin (Fisher Scientific/ Life Technologies, Illkirch, France). We read the results using a BioPlex-200 (BioRad, Marnes-la-Coquette, France) or MagPix (Luminex, Austin, TX, USA), expressing them as median fluorescence intensity (MFI) per 50 beads.

**Social science study**

Study A:

To complement the ecological study, we aimed to identify the daily activities carried out by the inhabitants according to their socio-demographic differences (age, gender, education, social status within the village, economic activities) that can trigger a spillover and the spread of the disease within the social and ecological environment of a rural village. This study included a structured questionnaire related to animal capture, human behaviour and subsistence activities, and local people's awareness, perceptions, and knowledge of bats (e.g. the month of the year when bats are present in local forests, the presence of bat roosts or breeding sites, bat hunting techniques and conditions of meat consumption, and beliefs or practices (e.g. medical virtues associated with bats, knowledge of bat-borne diseases). We interviewed volunteers (both female and male, older than 18 years) living in Inkanamongo. We selected people from a wide range of occupations (i.e., traditional, professional, and self-employed), and with a variety of interests in the forest and wildlife to get a diversity of opinions. We stopped interviewing when new data didn't add anything new to the understanding of the issue (data saturation). These interview sessions were combined with participant observations and informal discussions during the ecological team's research activities (such as collecting trapped animals, collecting specimens at the bushmeat market), and on the activities and movements of residents within and between villages in the study area. Data from the questionnaire were collected on paper forms in Lingala by an anthropologist from the DRC, translated in French and encoded in an Excel table for statistical analysis using R software. Notes and audio recordings from participant observations and informal discussions were transcribed in a Word document. We conducted a thematic analysis focusing on the understanding of the interactions between the ecosystem and the inhabitants of the village of Inkanamongo.

Study B**:**

To select the six villages, we used a list made by the Tshuapa Provincial Division of Humanitarian Affairs and Solidarity, which was responsible for psychological support to the affected communities during and after the 2014 outbreak. The survey was conducted door-to-door; one person per household (female or male, older than 18 years) was invited to answer our questions. Data was collected among 117 persons, among whom 29 were women (24.8%) and 88 were men (75.2%). **Table S3** shows the number of persons interviewed in every village. Data was collected in Lingala and/or Kimongo (locally spoken languages) and was simultaneously audio-recorded and registered manually through tablets in the REDCap software. A database (in an MS Excel file) was produced by REDCap. A team of research assistants based in Kinshasa translated the recordings into French. The French translations were automatically transcribed through the software SONIX. The information in the database was compared with the recorded, translated, and transcribed information. Relevant findings were contextualised through the participants' explanations (qualitative data) when possible.
In the six villages, we collected data about:

- Demographic characteristics of the respondent (age; gender; ethnicity; education level; religion; number of household members; social status (=role held in the community); primary activity; second and possible third occupation);
- Individuals' interactions with the forest (reasons for and frequency of entering the forest; periods of highest frequency of entering the forest; animals spotted; people (from the same or another village) accompanying the respondent to the forest; wild meat selling, buying,

consumption, and preparation for consumption; frequency of fetching water; people accompanying or dedicated to fetching water);

- Interactions between communities (individuals' travel destinations, reasons for and frequency of travelling, reasons for spending the night outside their village) and within the community (mass gatherings and estimated attendance);
- Individuals' health-seeking behaviours (under normal circumstances);
- Influence of rumours of an EVD outbreak on conducting daily activities.

**References supplementary material**

**List of supplementary tables**

**Table S1.** Overview of previous studies investigating Ebola in wild animals from 1976 to 2023.

**Table S2**. Primers used in this study.

**Table S3.** Number of persons interviewed in every village in Djera Sector.

**Table S4.** Reactivity to zero, one or two antigens of the same orthoebolavirus species depending on the history of contact with Ebola.

**Table S5.** Molecular identification of wild meat species from Inkanamongo hunters during the 2021 animal reservoir study.

**Table S6**. Animal taxa either reported as occurring in or around the Salonga National Park and their translation in local language (Lingala or Kimongo), their common name in French English and their scientific species name.

**List of supplementary figures**

**Figure S1.** Comparative analysis of IgG responses to Ebola virus antigen combinations in Inkanamongo (DR Congo) based on exposure.

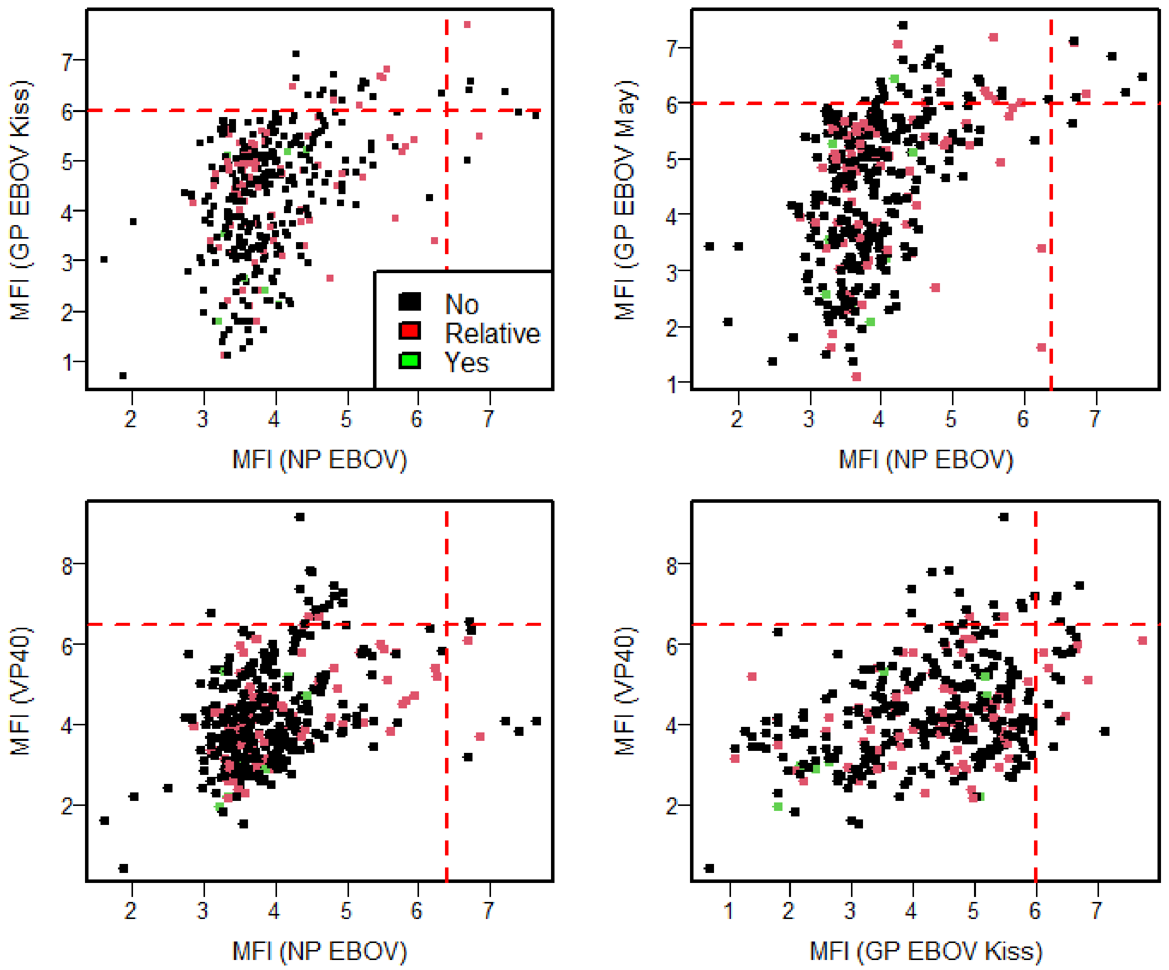


**Figure S1**. Comparative Analysis of IgG Responses to Ebola Virus Antigen Combinations in Inkanamongo (DR Congo) Based on Exposure. *IgG antibody titres in inhabitants of Inkanamongo (DR Congo), expressed as the log of Median Fluorescent Intensity (MFI) values measured by Luminex for four Ebola virus antigens: nucleocapsid protein (NP), glycoprotein from the Kissidougou strain (GP-Kiss), glycoprotein from the Mayaro strain (GP-May), and viral protein 40 (VP40). The data are presented as follows: GP-Kiss and NP (top left), GP-May and NP (top right), VP40 and NP (bottom left), and VP40 and GP-Kiss (bottom right). Each dot represents an individual participant. Participants are classified by Ebola virus disease (EVD) exposure: no infection (black), infected relative (red), and participant became infected (green). The red dotted line represents the antigen-specific cutoff on the Luminex assay.*
